# PANDA: A simple and affordable chamber system for measuring the whole-plant net CO_2_ flux

**DOI:** 10.1101/2025.06.02.657330

**Authors:** Philipp Schuler, Yuxin Li, Pierre Pittet, Patrick Favre, Mai-He Li, Yan-Li Zhang, Charlotte Grossiord

## Abstract

The carbon (C) balance of plants is the sum of all source and sink processes. However, due to methodological limitations, most studies focus predominantly on measurements of leaf-level assimilation and respiration, with less attention given to these processes in heterotrophic organs or the whole-plant level. As a result, knowledge of the whole-plant net C balance is scarce, limiting our understanding of the dynamics between C source and sink activities. Therefore, we developed an easily reproducible chamber system for continuous measurements of whole-plant net CO_2_ fluxes. We present the obtained dynamics of net CO_2_ fluxes of several C_3_ and CAM species, including germinating *Quercus robur*, over several days, as well as the whole-plant net CO_2_ flux temperature response of *Q. robur* seedlings, identifying the temperature thresholds at which they shift from a net CO_2_ sink to source. We show distinct diel patterns of net CO_2_ fluxes in C_3_ plants, likely driven by a dynamic diurnal up- and downregulation of sink activities in woody C_3_ plants. These patterns appear temperature-driven, suggesting a dynamic response of plant’s sink and source activity to environmental drivers. Our results highlight the importance of whole-plant C balance measurements for understanding plant responses to environmental conditions.

## Introduction

Determining net CO_2_ fluxes at the whole-plant level comes with difficulties, since it integrates several parallel autotrophic and heterotrophic fluxes over the whole plant body, including leaves, branches, and roots. Most commonly, gas-exchange measurements are conducted at the leaf-level to quantify net assimilative CO_2_ fixation (A_net_) and respiratory (R) CO_2_ release. In contrast, fewer studies address fluxes from other organs, such as branches, stems (Han et al., 2017; Diao et al., 2020; Zhang et al., 2025), and roots (e.g., Atkin et al., 2000; Kuzyakov and Larionova, 2006). These organs, especially roots, act as major carbon (C) sinks and contribute significantly to the plant’s overall C balance by consuming up to more than 50% of all assimilates (Lambers et al., 2002; Colombi et al., 2021), and it is known that the root- and stem-level metabolic responses to temperature can have a significant impact to whole plant level C relations (Atkin et al., 2000; Wang and Hoch, 2022). Moreover, twig and stem CO_2_ fixation can also significantly contribute to the whole plant C balance, reassimilatint 7 to 123% of respired CO_2_ in non-succulent species (Ávila et al., 2014). Still, there are methodological difficulties in measuring the net CO_2_ flux of a particular piece of twig or stem, for example, stem photosynthesis can occur in plant species with and without stomata on these organs (Ávila et al., 2014), making measurements difficult to interpret and hard to compare. Furthermore, CO_2_ is being redistributed within the plant xylem by the sap stream (Teskey et al., 2008; Grossiord et al., 2012; Trumbore et al., 2013), causing strong reduction of the measured CO_2_ efflux of up to 40 % under high xylem sap velocity (Gansert and Burgdorf, 2005). Moreover, diel patterns of stem CO_2_ efflux differs between phylogenetic groups such as monocots, cycads, as well as angiosperm and gymnosperm trees (Marler and Lindström, 2020), making an accurate comparable temporal tracking and quantification of tissue-specific measurements to the whole plant level challenging.

Measuring whole-plant net CO_2_ fluxes remains challenging due to the technical difficulties of enclosing entire plants, accurately capturing integrated gas exchange over time, and simultaneously excluding soil respiration, therefore, only few studies have been able to measure whole-plant net CO_2_ fluxes (Hand, 1973; Dutton et al., 1988; Zhao et al., 2013; Brauner et al., 2014, 2018; Patono et al., 2023). This methodological gap limits our ability to fully understand how entire plants respond to environmental stressors under realistic conditions. One study that successfully measured whole plant net CO_2_ fluxes demonstrated that the carbon balance of *Thuja occidentalis* L. becomes negative - reflecting respiratory losses exceed photosynthetic gains - at temperatures above 35°C and mild drought (Zhao et al., 2013). This study highlights the high potential of such measurements for understanding plant responses to climatic stress and for predicting future shifts in the global C cycle. Recent systems developed for tracking the net CO_2_ flux in larger plants (Patono et al., 2023), while valuable, are often large, technically complex, and relatively difficult to reproduce, making them impractical for routine use in standard growth facilities or climate chambers. Smaller-scale systems suitable for controlled environments frequently measure only the aboveground biomass (Kölling et al., 2015; Salvatori et al., 2021), which strongly bias estimates of whole plant net CO_2_ balance - especially in larger plants such as trees, where root biomass can be comparable to that of the canopy (Perry, 1982; Harris, 1992). However, existing small systems to capture whole plant net CO_2_ flux are designed to measure small plants such as *Arabidopsis thaliana* (L.) Heynh.(Brauner et al., 2014, 2018), and therefore are not suitable for larger plant species such as woody plants.

Nevertheless, whole-plant C exchange measurements offer a key advantage in addressing current hot topics in plant science by capturing integrated responses that are missed by regular tissue-level measurements or ecosystem-scale approaches like eddy covariance, which are showing discrepancies in their evaluation of the whole-plant C dynamics (Campioli et al., 2016). For instance, it is crucial to isolate how the C balance of whole plants is affected by the ongoing exacerbation of heatwaves (Meehl and Tebaldi, 2004; Perkins-Kirkpatrick and Lewis, 2020), as these events can differentially impact photosynthesis, respiration, as well as C allocation across tissues (Birami et al., 2018; Werner et al., 2020). For instance, stem photosynthesis can buffer against the sharp decline in leaf photosynthesis during heatwaves (Ávila-Lovera et al., 2024) - effects that are only fully captured when measured at the whole-plant scale. Furthermore, direct measurements of whole plant net CO_2_ fluxes could have the potential for a faster selection of heat-resistant genotypes in plant breeding, both in agriculture and forestry. Finally, Earth System Models are used to simulate the future pace and extent of CO₂ increases and climate change (Prinn, 2013), integrating multiple subcomponents such as plant functional types, vegetation dynamics, C fluxes, as well as their response to climate on high temporal resolution. Therefore, an accurate knowledge of whole plant net CO_2_ fluxes of different functional types responds to changes in temperature, how they change during plant development, and on a diel course, is essential for accurate predictions of the future trajectories of the global atmospheric CO_2_ concentration.

To address these gaps, we developed an easy-to-replicate and low-cost chamber system (**P**lant**A**ssimilatio**ND**yn**A**mics (PANDA) that can be attached to any gas exchange system capable of measuring and regulating CO_2_ concentrations, such as the WALZ GFS-3000 (Heinz Walz GmbH, 91090 Effeltrich, Germany) or the Li-6800 system (LI-COR Environmental, Lincoln, NE 68504, United States). To validate this system while simultaneously addressing a key uncertainty in whole-plant carbon fluxes, namely how daytime, growth and temperature influence the whole plant net CO₂ balance, we designed four experiments:

- To assess acclimation effects to the system, we first continuously measured the whole plant net CO_2_ balance of three C_3_ (*Brachychiton acerifolius* (A.Cunn. ex G.Don) F.Muell., *Trachycarpus fortunei* (Hook.) H.Wendl., *Coffea arabica* L.) and three CAM species (*Aloe arborescens* Mill., *Opuntia* Mill., *Phalaenopsis* Blume), the latter which exhibit distinct and well-studied diel changes in their net CO_2_ flux (Osmond et al., 1979), over two to three diel cycles. This experiment allowed us to test whether the measurements of a plant remains constant for several days of normal day/night cycles.
- In a second experiment, we monitored the whole-plant net CO_2_ flux during several days of constantly light or dark conditions to investigate whether the system can be used to identify circadian cycles of the net CO_2_ balance independent of light availability.
- To provide first ideas for potential plant physiological investigations with the system, we monitored the net CO_2_ balance of germinating *Quercus robur* L. acorns over one month.
- Finally, we subjected the *Q. robur* seedlings twice to increasing temperatures of about 27, 34, and 43°C. Once after the first seedling growth fully finished, and once after the plants produced a second flush. With this, we could investigate potential changes of the response due to changes in ratios of autotrophic to heterotrophic tissues between the curves after the first and second flush.

## Materials and Methods

### Chamber design and use

The **P**lant**A**ssimilatio**ND**yn**A**mics (PANDA) chamber (Fig. 1) consists of (Fig.1b, c) a 30 × 30 × 1 cm bottom plate (1) that features a circular recess (2; 24.5–25.5 cm diameter) accommodating air inlet/outlet ports (AI, AO) from and to the gas analyzing system (we used a GFS-3000 Gas-Exchange System, Heinz Walz GmbH, Germany), a water inlet (WI) for watering, where a watering hose is connected on both sides of the plate, and a cable gland (CG). Air and water inlet ports were connected with silicone tubes on both sides of the plate. Metal feet (F) measuring 8 cm are fixed at each corner. The internal fans and sensor connect to and are controlled by a Raspberry Pi 5 (Raspberry Pi Foundation, England) positioned outside the chamber, with live data displayed on an attached screen. Inside, two axial fans (V; Sunon MF60151V2-1000U-A99, Sunon Fans, Taiwan) and a temperature/humidity sensor (TH; Adafruit Sensirion SHT45 Precision Temperature Humidity Sensor, Adafruit, USA) are present. The chamber (a) is closed with an 80 cm tall, 25 cm diameter glass cylinder (C; Sandra Rich GMBH, Germany). To prevent condensation water from reaching the gas analyzer, a water trap (WT) had to be added to the air outlet tube outside the chamber, approximately 1m away from both the chamber and gas analyzer, with the water trap representing the lowest position of the whole system. It must be noted that the PANDA measuring chamber was not designed to regulate air temperature, relative humidity, or light but rather to serve an enclosed chamber to measure gas exchange fluxes. The temperature and light must be defined either by natural light and ambient temperature variation (e.g., by using the system outdoors) or actively being regulated with additional devices, such as heating units, LED light sources, or by placing the system inside a climate chamber. Due to the high evapotranspiration, the relative humidity (RH) raises rapidly (usually >80%), especially in well-irrigated plants, when connected to a standard gas analyzing system with a comparable low flow rate, such as the WALZ GFS-3000, which was used in this study. Hence, for researchers aiming to tightly control RH, we recommend either increasing the system’s flow rate, which must use a more powerful control unit than the here used WALZ GFS-3000, or by integrating a bypass desiccant system.

**Figure 1:**
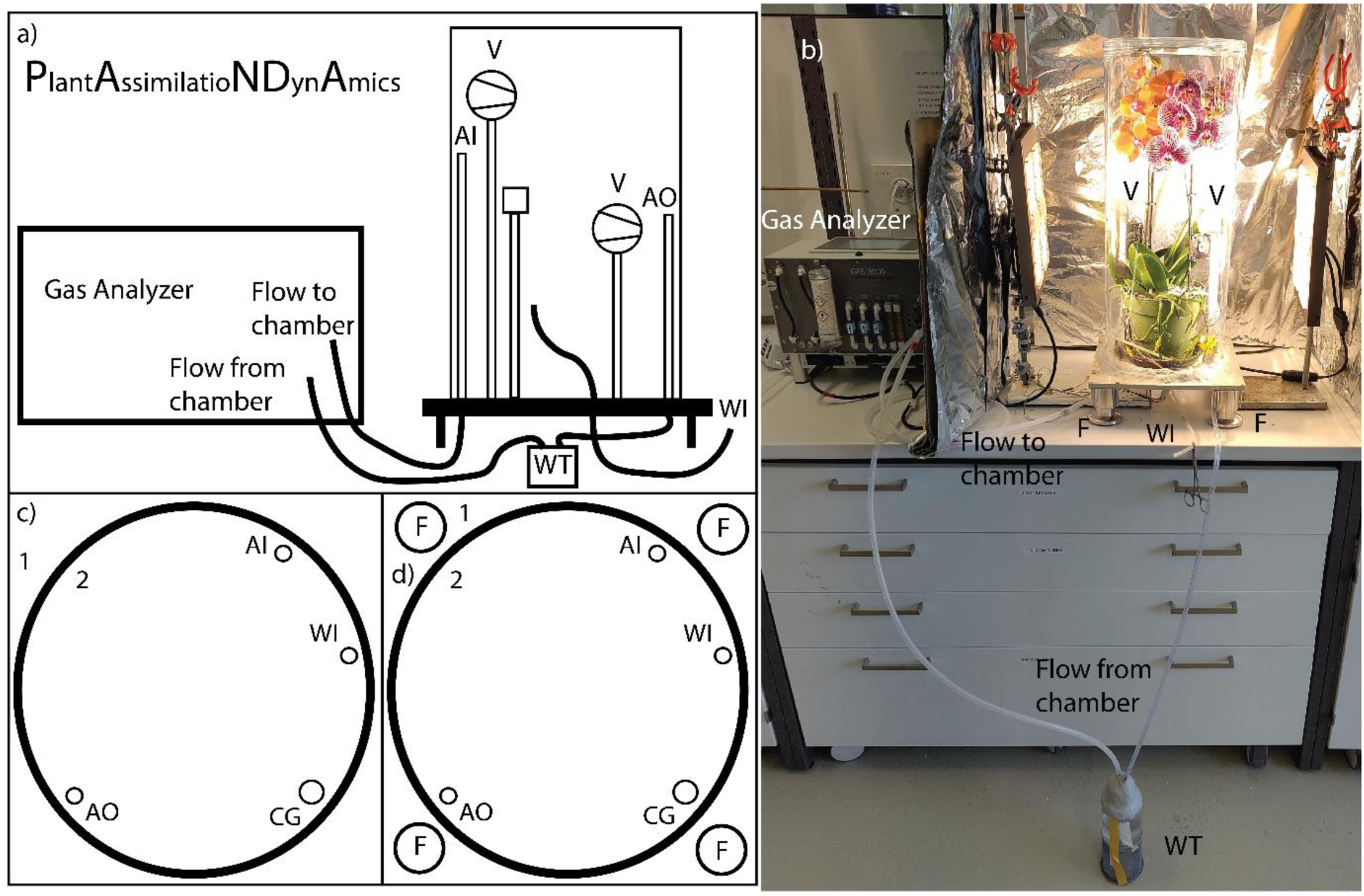
The (a, b) PlantAssimilatioNDynAmics (PANDA) chamber consists of (c, d) a bottom plate (1; 30 x 30 x 1 cm) with a circular recess (2) with an inner diameter of 24.5 cm and an outer diameter of 25.5 cm milled into it. Inside the recess, an air inlet port (AI), an air outlet port (AO), a water inlet port (WI), and a cable gland (CG) are inserted. On each corner of the plate, 8 cm high metal feet (F) are attached. Inside the chamber (a), two fans (V; Sunon MF60151V2-1000U-A99 axial fan 12 V/DC) as well as a temperature and humidity sensor (TH; Adafruit Sensirion SHT45 Precision Temperature Humidity Sensor). The fans and the sensor are connected and controlled, as well as the data stored on a Raspberry Pi 5, which is located outside the chamber, the latter of which was attached to a screen for visually tracking chamber conditions (not shown in the scheme). To prevent condensation water from reaching the gas analyzer, a water trap (WT) is installed in the air outlet tubing, approximately 1m away from both the chamber and gas analyzer, with the water trap representing the lowest position of the whole system. The chamber is sealed by using an 80 cm high glass cylinder with a diameter of 25 cm (C; Glass vase CYLI clear cylindrical, Sandra Rich).

Depending on the specific requirements of the planned experiments, one can add more access ports to the bottom plate of the chamber. For instance, we recommend adding an outlet port for condensation water, which can accumulate at the bottom of the chamber, especially during long-term measurements. Furthermore, the chamber size can also be adjusted by using different sizes of transparent (glass) cylinders or similar. The main limitation is the maximum flow rate of the analyzer unit, which limits the amount of CO_2_ that can be added. Since we were working with two light sources on two sides of the chamber (Fig. 1b), we installed four panels covered in aluminium foil on each side of the chamber (only shown three of four in Fig. 1b) to provide equal high light availability around the plants.

### Calculation of the net CO_2_ flux

The net CO_2_ flux in µmol s^-1^ was calculated as:

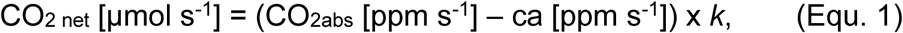

with CO_2abs_ being the CO_2_ concentration delivered to the chamber, ca is the CO_2_ concentration that is coming from the chamber, and k is the conversion factor to convert the measured CO_2_ flux from ppm into µmol, which is calculated as follows:

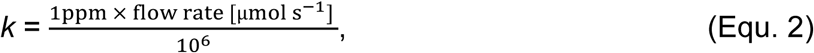

leading to a k of 0.0009 μmol CO_2_ s^−1^ ppm^−1^ at a flow rate of 900 µmol s⁻¹. Due to the large chamber volume, we choose the maximum flow rate possible for the system. To convert the resulting µmol CO_2_ s^-1^ into µmol glucose s^-1^, one can divide µmol CO_2_ s^-1^ by 6, representing the stoichiometric C ratio of 1 µmol CO₂ to 1 µmol glucose.

### Calculation of the cumulative CO_2_ and C flux

The cumulative CO_2_ and C flux was calculated as:

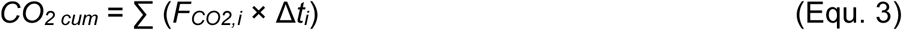

where F_CO2,*i*_ is the net CO₂ flux at time point *i* (µmol CO₂ s⁻¹), and Δ*ti* is the time interval between consecutive measurements (seconds). The summation is done over all time intervals from the start to the end of the period of interest. If the time intervals are equal (e.g., measurements every minute), the formula simplifies to:

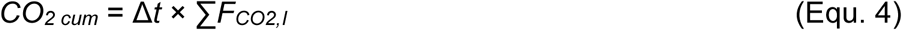

Example Calculation:

If you have flux measurements every minute (60 seconds) and the net CO₂ flux at three consecutive time points is 0.5, 0.7, and 0.6 µmol s⁻¹, the cumulative flux after 3 minutes would be:

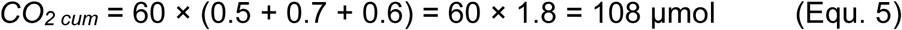

and to finally convert the cumulative C balance into gravimetric units, we divide the resulting μmol (CO_2_) by 6 to convert it into μmol (Glucose), and multiply this by the molar mass of glucose:

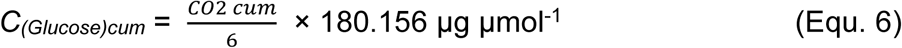

### Carbon-free substrate for plant growth

The substrate used for the net CO_2_ flux measurements is pure C-free perlite (Pflanzen Perlit, grain size 2-6mm, Otto Hauenstein Samen AG, Switzerland). The advantage of pure C-free perlite is that soil respiration is excluded from the measurement, as there is no organic matter or microbial C source to contribute to CO_2_ emissions. Furthermore, since perlite possesses high porosity, it ensures rapid CO₂ exchange between the substrate and the chamber air space. Another advantage of using this pure white perlite is that it is easy to see contaminations such as algae or other microorganisms growing at its surface, something of great importance, especially during long-term measurements. As perlite lacks nutrients, they need to be added externally when plants are cultivated for a longer period of time, however, this also ensures that specific nutrient concentrations can also be precisely adjusted to the experimental needs. The high pore volume is providing well-aerated conditions at the root level but a relatively limited water-holding capacity (Jackson, 1974) and therefore requires careful watering, which may not reflect the soil conditions certain plants typically experience in natural environments. Hence, root development and function might differ from those in more structured or biologically active soils, potentially influencing whole-plant physiological responses and C dynamics. Thus, it might be beneficial to grow plants in a more natural soil prior to placing them in the PANDA chamber, using C-free perlite only during the experimental stage. However, an optical check of the root development of the tested oak seedlings (Fig. S1) revealed no abnormalities, indicating normal root growth under the tested conditions.

### Plant cultivation prior to measurements and plant positioning in the chamber

Plants are grown in perlite for at least 1 month to ensure the roots are acclimated to the porous growing material and the experimental light conditions. Plants were fertilized once per week with liquid inorganic fertilizer (Blumen-Dünger 1l (N-P-K 4-3-3), terrasan Schweiz GmbH, 8034 Zürich, Switzerland). The room temperature was 22-24°C, and the RH was ∼50-60%. Plants were grown under a 100W LED plant light (grow-led-pflanzenlampen.ch, Senn Services, Switzerland) with a 12-hour day/night cycle. To ensure that no organic C from root exudates or shedded root cells was contaminating the substrate, two days before the start of the experiment, the plants were carefully removed from the pots, the old perlite was carefully washed away from the roots, and the plants were repotted in new perlite. Therefore, any contribution of soil respiration to the measurements could be excluded.

On the day the measurements started, plants were positioned with a horticultural saucer in the center of the bottom plate. The inner end of the watering hose was fixed below the pot, and the outer end of the watering hose was closed with a hemostatic clamp. After that, waterproof multi-purpose sealant (TEROSON RB IX, Henkel AG, Germany) was added by forming a 1 cm high and 2 cm wide wall on top of the circular recess, the glass cylinder was carefully positioned on the sealant, and the cylinder was tightly connected by applying gentle pressure on the top surface by weight. On two sides of the glass cylinder at a distance of 15cm, a 100W LED plant light (grow-led-pflanzenlampen.ch, Senn Services, Switzerland) was installed, running as well in a 12-hour day/night cycle. The photosynthetic active radiation, measured with the LI-6800 in the center of the bottom plate, was 1250 µmol m^-2^ s^-1^ when measured in the direction of either plant light. The temperature inside the chamber increased from an average of 22.1°C at night to an average of 31.8°C during the day (Fig. S2) due to the heat generated by the light. Due to the high evapotranspiration rate, the relative humidity (RH) was above 90%, so the vapor pressure deficit (VPD) remained below 1 kPa (Fig. S2). This temperature and RH regime was the same for all experiments in the next subsection, except for the temperature response curves of experiment 4.

### Continuous measurement of the whole plant net CO_2_ flux

The measurements were done by running an auto-program with the GFS-3000 system, which regulates the CO_2_ concentration inside the chamber (ca ppm) at 420 ppm with a total flowrate of 900 µmol s^-1^. Inflowing (CO_2abs_), outflowing (ca ppm), as well as the delta between the two (dCO_2_MP), were recorded every 2 minutes. Plants were watered by filling the horticultural saucer using a squeeze bottle via the water inlet port.

To evaluate the performance of the system, we performed four experiments:

### Experiment 1: Diurnal variation in whole-plant CO_2_ exchange in C_3_ and CAM plants

To test the sensitivity of the system and whether the results of the daily measurements are comparable between different days of measuring due to acclimation effects, we tracked the net CO₂ flux of single plants of three C_3_ (*Brachychiton acerifolius* (A.Cunn. ex G.Don) F.Muell., *Trachycarpus fortunei* (Hook.) H.Wendl., *Coffea arabica* L.) and three CAM species (*Aloe arborescens* Mill., *Opuntia* Mill., *Phalaenopsis* Blume) over two to three diel cycles. Woody C_3_ plants were included to investigate whether diel patterns could be observed in their net CO_2_ flux, as has been observed in C_3_ forbs (Resco De Dios et al., 2016), and CAM species, which are well known for their four diel oscillatory cycles of CAM photosynthesis (Osmond et al., 1979), were included to evaluate the ability of the PANDA chamber to identify diel pattern. To compare the whole-plant net CO_2_ flux with leaf-level net assimilation (A_net_ at PAR = 1500, RH = 60%, T_air_ = 30°C), leaf gas exchange was tracked with a Li-6800 over several hours of three leaves per plant.

### Experiment 2: Endogenous cycles in C_3_ plants

Additionally, we measured the net CO_2_ flux of the C_3_ plants under three to four days of continuous light or dark conditions to investigate potential endogenous rhythms (Lüttge and Hertel, 2009) in their source- and sink dynamics. Indeed, many plants exhibit cyclical patterns of CO₂ uptake and release that follow a ∼24-hour rhythm. This suggests that photosynthesis and respiration are partially regulated by internal clocks, not just light availability (Resco De Dios et al., 2016). Understanding the extent of endogenous rhythms in CO_2_ fluxes is important for accurate interpretation of gas exchange data, for the design of experiments, and for understanding and modeling plant C balance and growth.

### Experiment 3: Net CO_2_ flux of establishing seedlings prior and during drought stress

To evaluate whether the system can accurately track the C balance of developing plants over a longer period, we continuously tracked of the net CO_2_ balance of germinating *Quercus robur* L. acorns. For this, we carefully peeled 15 slightly germinating (with initial root growth) acorns, weighed their initial weight, and planted 12 of them into fresh perlite in the same pot, and determined the initial dry mass of the remaining 3 acorns by drying them in the oven at 60°C for 7 days. The planted acorns and the pot have been placed inside the chamber, and the net CO_2_ flux was continuously tracked over a whole month. Groups of three seedlings were carefully harvested after 9, 13, and 30 days to track the change in dry biomass of leaves, shoots, and roots after drying them in the oven at 60°C for 7 days. Finally, to study the impact of a mild drought on the net CO_2_ flux of the seedlings, the plants were subjected to a mild drought by stopping the irrigation during the last four days of the germination experiments.

### Experiment 4: Shifts in whole-plant C balance with rising temperatures

After the seedlings of experiment 3 finished their first growing cycle and had fully mature leaves, the three remaining plants were carefully taken out of the pot and the spent perlite was washed away. The plants were repotted in new perlite, again all plants in a single pot. Subsequently, the first of two temperature response experiments of whole plant net CO_2_ flux was conducted (Round 1). For this, the plants were subjected to increasing target air temperatures of ∼27, ∼34, and ∼43°C over three days (i.e., changed temperature each day). The actual temperature values are displayed in Fig. S3. To decrease the air temperature in the system during the light hours of the first day, the temperature input of the light to the chamber surface was decreased by cooling the glass cylinder to about 27°C with two external desk fans. On the second day, the temperature input by the light was not altered, leading to a T_air_ of about 34°C, while on the third day, T_air_ was increased by carefully heating the system with an external heating vent (Trisa Heizlüfter Heat & Chill 2’000W, Trisa Electronics AG, Switzerland) to about 43°C.

After this first temperature response experiment, the plants were flushed anew, and the plants grew inside the chamber until the leaves and shoots matured. While the new leaves and shoots were fully grown, the color of the new leaves was still slightly paler green compared to the ones of the first flush, indicating that their chlorophyll concentration did not reach the maximum. However, since the maximum CO_2_ input into the system would have been surpassed, we continued with the second temperature response of the same temperature range (round 2) to compare the resulting response curves of the two plant sizes.

Leaf area and biomass were determined by harvesting three seedlings after both rounds of temperature responses. Leaves were scanned (CanoScan LiDE 300, Canon, Tokyo, Japan), and the total leaf area was analyzed with ImageJ (Schneider et al., 2012). The dry weight was measured (NewlassicMF, MS105DU, Mettler Toledo, Switzerland) after drying them in the oven at 60°C for 7 days.

## Results and Discussion

### Experiment 1: Diurnal variation in whole-plant CO_2_ exchange in C_3_ and CAM plants

The net CO_2_ flux measured with the PANDA chamber (Fig. 2) showed robust tracking of the diel CO_2_ fluxes over several days, which was reproducible between the different days, both in C_3_ (Fig. 2a) and CAM (Fig. 2b) plants, thereby demonstrating the reliability and functionality of the system for capturing consistent whole-plant gas exchange dynamics.

**Figure 2:**
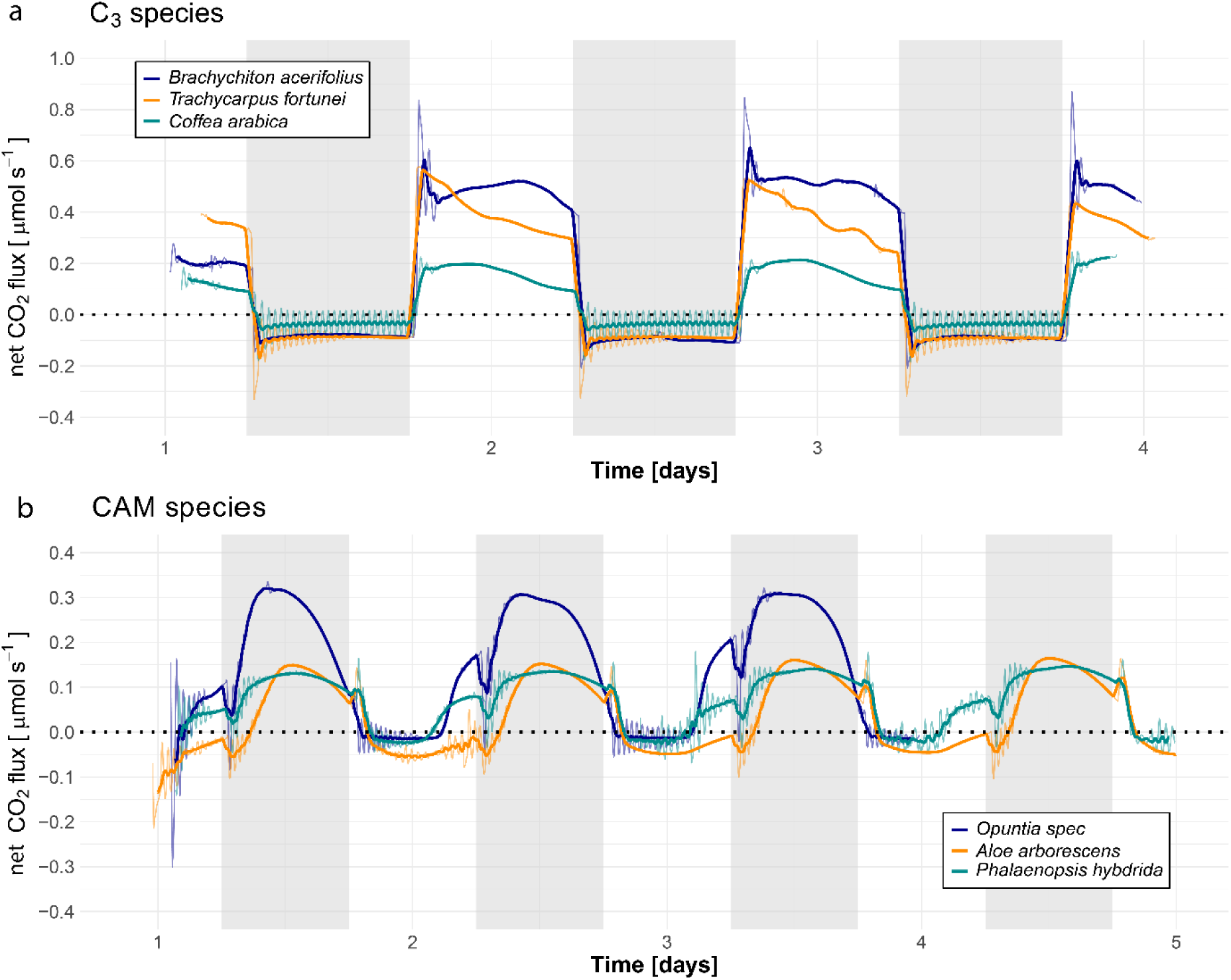
Diel dynamics of the net CO_2_ flux of (a) the three measured C_3_ and (b) CAM species over three to four days (n = 1 per species). Nighttime hours are shown in grey. The bold lines show the running means (10 measurements before and 10 measurements after each point, with one measurement every 2 minutes), and the fine lines show the corresponding measurements.

All C_3_ plants showed little variability in their CO_2_ fluxes during the nighttime (Fig. 2a). However, their positive daytime fluxes were (i.e., indicating higher C uptake than release) highest at the beginning of the day, and showed species-specific trends, usually having the lowest net CO_2_ balance just before the light turned off. Interestingly, the net CO_2_ flux of the two dicotyledons *B. acerifolius* and *C. arabica* spiked, the former also with the overall most positive net CO_2_ flux, at the beginning of the day, followed by a slow and steady increase after they reached a plateau, before declining steadily for the remainder of the day. However, the midday peak was less consistent between the two days of measurement in *B. acerifolius*. The smaller spike in *C. arabica* compared to *B. acerifolius* could be explained by its relatively low photosynthetic temperature optimum of 24°C (Nunes et al., 1968; Kumar and Tieszen, 1980; DaMatta et al., 2018), consistent with its origin in the cooler montane rainforests of Ethiopia (Kufa and Burkhardt, 2011). In contrast, *B. acerifolius*, native to tropical lowland forests (“CSIRO - Australian Tropical Rainforest Plants,” 2020) is better adapted to temperatures above 30°C. *T. fortunei*, a monocot palm, also showed a peak in net CO_2_ flux at the beginning of the day, followed by a steady decline throughout the daylight hours. This trend may be partly explained by a continuous increase in meristem temperature, which is known to increase the growth rate in monocots (Ben-Haj-Salah and Tardieu, 1995) and contribute to a steady increase in growth respiration. At the leaf-level, the net assimilation rate in the C_3_ species (Fig. S3) were less consistent under conditions of 30°C and 60% RH over several hours, suggesting that the daytime flux variability may result from complex interactions between C source and sink activities and the environment, such as an increased sink activity due to slowly increasing soil and shoot temperatures (Wang et al., 2006) and an increased availability of fresh assimilates during daytime hours increasing respiration (Lai et al., 2016; Brauner et al., 2018). At night, all three species exhibited peak respiration at the start of the dark period, likely caused by residual heat and substrate availability following light-off (Frantz, 2004). Thereafter, whole plant net CO_2_ flux remained consistently negative and stable, indicating that plant C dynamics are more variable during the day than at night, which is in contrast to earlier studies in forbs where distinct patterns during the nighttime were found (Gessler et al., 2017). This might either indicate that forbs regulate their respiration differently during the night, or that the nighttime C fluxes are coordinated on the whole-plant level.

Measurements of the CAM plants (Fig. 2b) clearly captured all four classical phases of CAM photosynthesis, as described by (Osmond, 1978). In phase I, during the night, stomata open to allow CO₂ uptake. In phase II, as the light switches on, stomata begin to close to prevent excessive water loss. However, during this transitional phase, both PEP carboxylase and Rubisco are active, allowing simultaneous CO₂ fixation via the CAM and C_3_ pathways. This dual activity results in small, visible spike in net CO_2_ flux at the beginning of the day (not observed in *Opuntia*). In phase III, with stomata fully closed during most of the day, water loss is minimized, thus only respiratory (i.e., negative) CO_2_ fluxes of heterotrophic tissues such as roots can be measured. Finally, in phase IV, in the late afternoon and absence of water stress, stomata reopen briefly, allowing additional CO₂ uptake via the C_3_ pathway. This is reflected in the observed in net CO_2_ flux late in the day. Following the transition to darkness, a sharp decline in net CO_2_ flux can be observed, marking the metabolic shift from C_3_ back to CAM pathway.

Together, these results highlight distinct diurnal CO₂ exchange patterns in C_3_ and CAM plants, emphasizing the PANDA chamber’s utility in resolving species- and pathway-specific gas exchange dynamics over time, such as cryptic or low level CAM photosynthesis (Winter and Holtum, 2015; Surridge, 2019; Winter et al., 2019).

### Experiment 2: Circadian cycles in C_3_ plants

Species-specific responses emerged when the C_3_ plants were exposed to extended periods of constant light or darkness (Fig. 3). In *B. acerifolius* (Fig. 3a), net CO_2_ flux stabilized over the three days of constant light, with the highest variability occurring on the first day. Under constant darkness, its net CO_2_ flux steadily declined, becoming increasingly negative over time. In contrast, *T. fortunei* (Fig. 3b) displayed pronounced fluctuations in net CO_2_ flux throughout the light period, with multiple sharp increases and decreases. This pattern might indicate an inherent cyclic growth rhythm in this monocot palm species, which becomes disrupted under prolonged light exposure. In *C. arabica* (Fig. 3c), the initially positive net CO_2_ flux during constant light gradually declined over the three days, exhibiting a characteristic wavelike or cyclic pattern. This decline could reflect either a gradual reduction in photosynthetic activity due to sustained exposure to high light and temperatures above 30°C, or a progressive increase in sink demand driven by continuous assimilate supply. The latter interpretation suggests a relatively stable source-sink coordination, potentially governed by recurring episodes of elevated sink activity. Under constant darkness, net CO_2_ flux in both *B. acerifolius* and *C. arabica* became progressively less negative, possibly indicating a depletion of non-structural carbohydrates (Weber et al., 2018) and a corresponding decline in sink activity (Sevanto et al., 2014; Collins et al., 2021). In contrast, *T. fortunei* showed no such trend, likely due to the mobilization of substantial non-structural carbohydrate reserves inside its stem (Tomlinson, 1990). However, the absence of clear patterns in respiratory nighttime CO_2_ flux is, as experiment 1, in contrast to earlier studies in forbs, where distinct patterns during the nighttime were found (Gessler et al., 2017).

**Figure 3:**
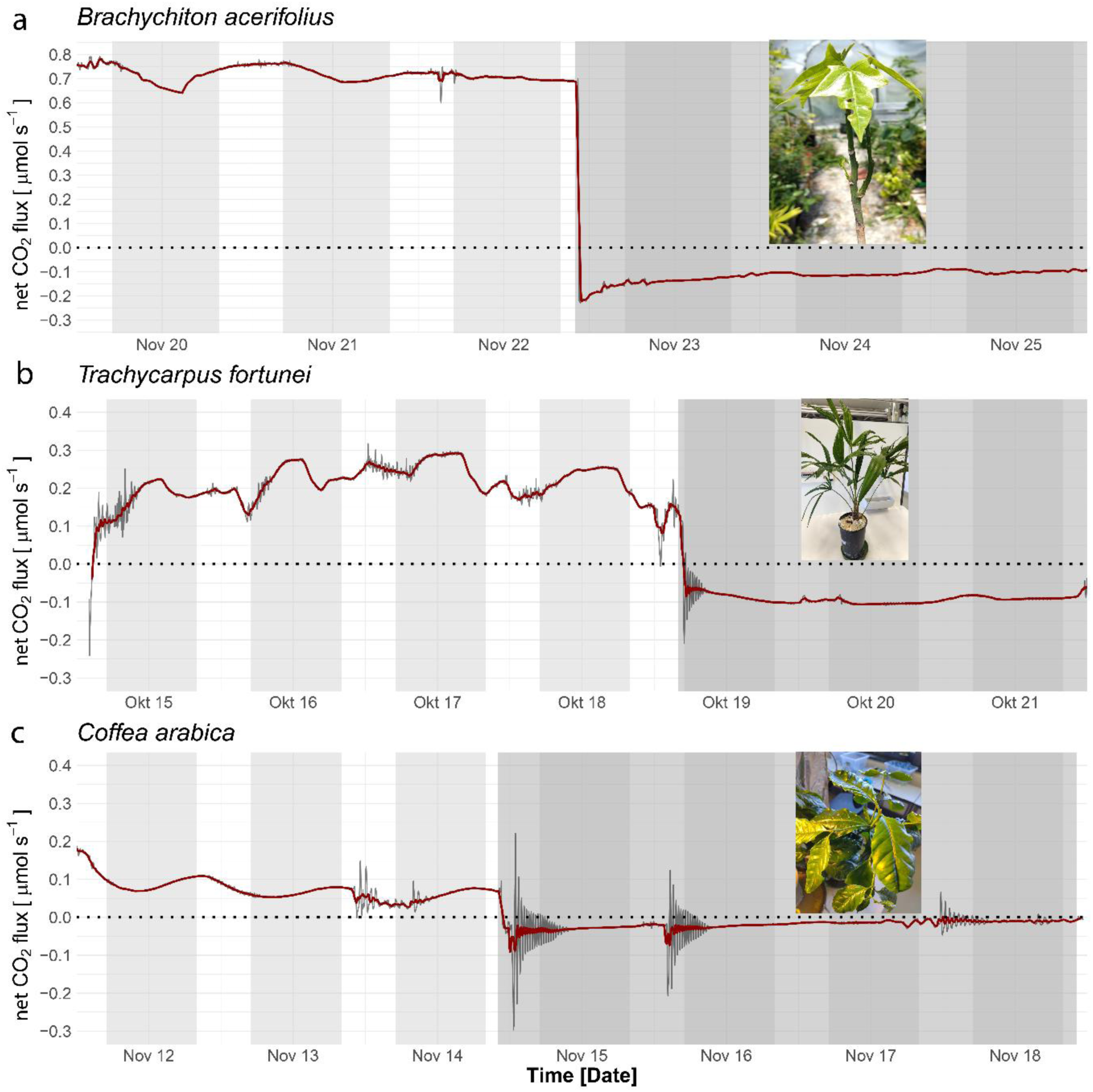
The net CO_2_ flux of the three tested C_3_ species (n = 1 per species) when exposed to three to four consecutive days of constant light and darkness. The lighter grey marks indicate the hypothetical nighttime hours, while the larger, darker grey area on the right indicates the actual time in darkness. The red lines show the running means (10 measurements before and 10 measurements after each point, with one measurement every 2 minutes), and the grey lines show the measured values.

These findings demonstrate the PANDA system’s capacity to resolve subtle, species-specific endogenous rhythms in CO₂ exchange, highlighting differences in source–sink coordination and carbohydrate storage strategies among C_3_ plants under constant environmental conditions. Future work should investigate the circadian regulation of the whole-plant net CO_2_ flux between different phylogenetic groups such as monocotyledons and dicotyledons, or angiosperms and gymnosperms, as well as different life forms, such as grasses, forbs, palms, and trees.

### Experiment 3: The net CO_2_ flux over one month of germinating *Quercus robur* acorns

Tracking acorn germination with the PANDA system (Fig. 4) demonstrated its ability to monitor net CO_2_ flux in growing plants continuously and accurately over more than a month. Mean daytime temperature and RH were 31.8°C and 96.6% (± 0.6°C and ± 2.6%, respectively), while nighttime values were 22.1°C and 100% (±0.4°C and ±0%). Net CO_2_ flux (µmol s^-1^ acorn^-1^) (Fig. 4a) captured the emergence of diel patterns from germination on February 22^nd^ to the fully developed seedling on March 19^th^, including the onset of a short drought treatment on March 21^st^. Seedling development at selected time point is illustrated in Fig. 5.

**Figure 4:**
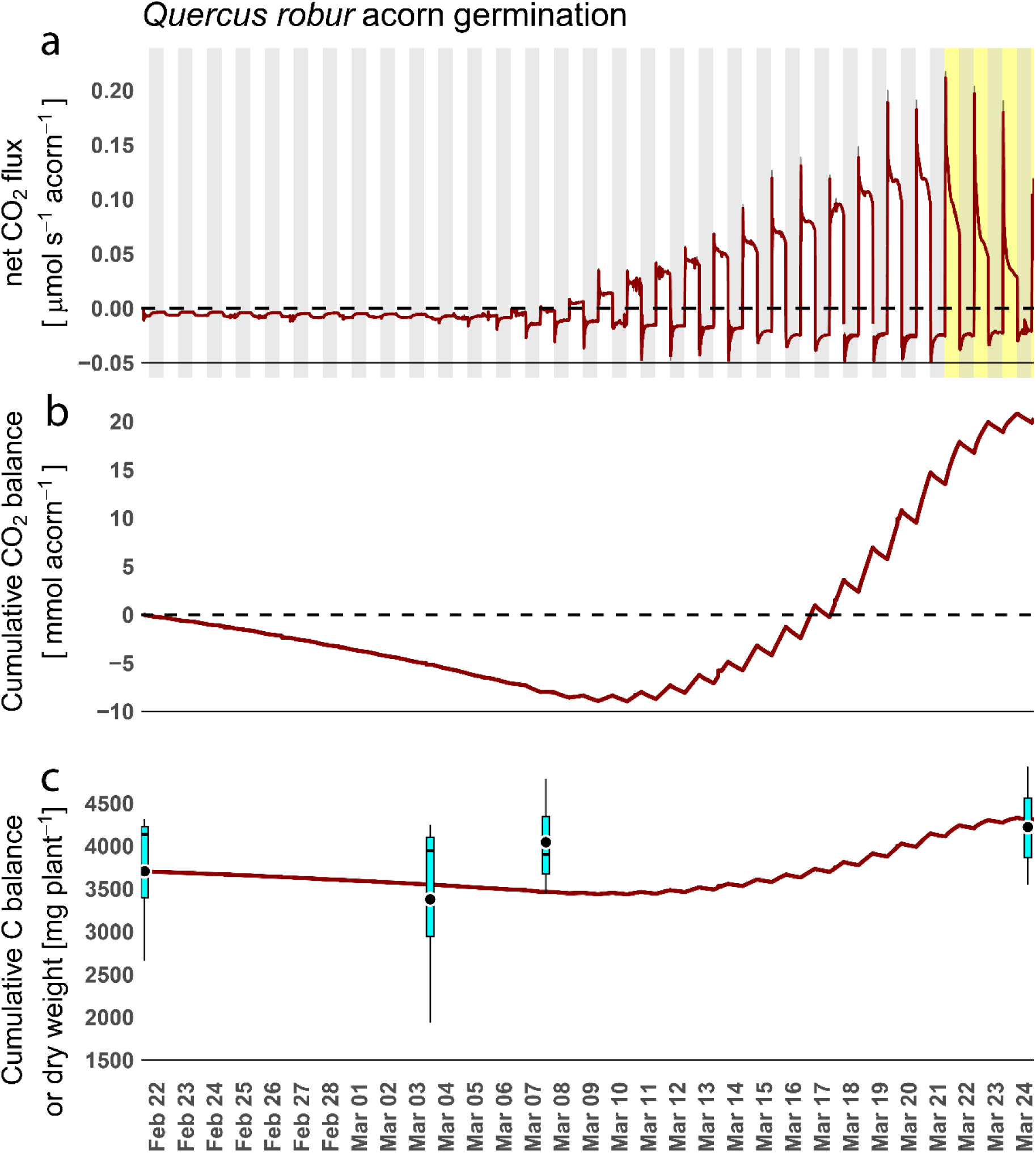
The (a) net CO_2_ flux in µmol s^-1^ acorn^-1^ per plant (12 replicates until March 3, 9 until March 7, and 6 until March 24) of germinating acorns of *Quercus robur* over one month, (b) the cumulative CO_2_ balance in mmol acorn^-1^ during this period, and (c) the calculated resulting change in biomass based as cumulative C balance in mg per plant calculated by the measured CO_2_ fluxes (red line) as well as the dry mass measured during the subsequent harvesting of three seedlings for the corresponding date (light blue boxplots). In (a), the light grey areas indicate the nighttime hours, and the yellow area indicates the days with reduced water access. The bold line in (a) shows the running means (10 measurements before and 10 measurements after each point, one measurement every 2 minutes), and in (b) and (c), the bold red line shows the actual values. In (c), the central horizontal line within each box represents the median of 3 harvested plants, while the box edges correspond to the 25^th^ and 75^th^ percentiles. Whiskers extend to data points within 1.5 × interquartile range. The black circle indicates the mean value for each harvest date.

**Figure 5:**
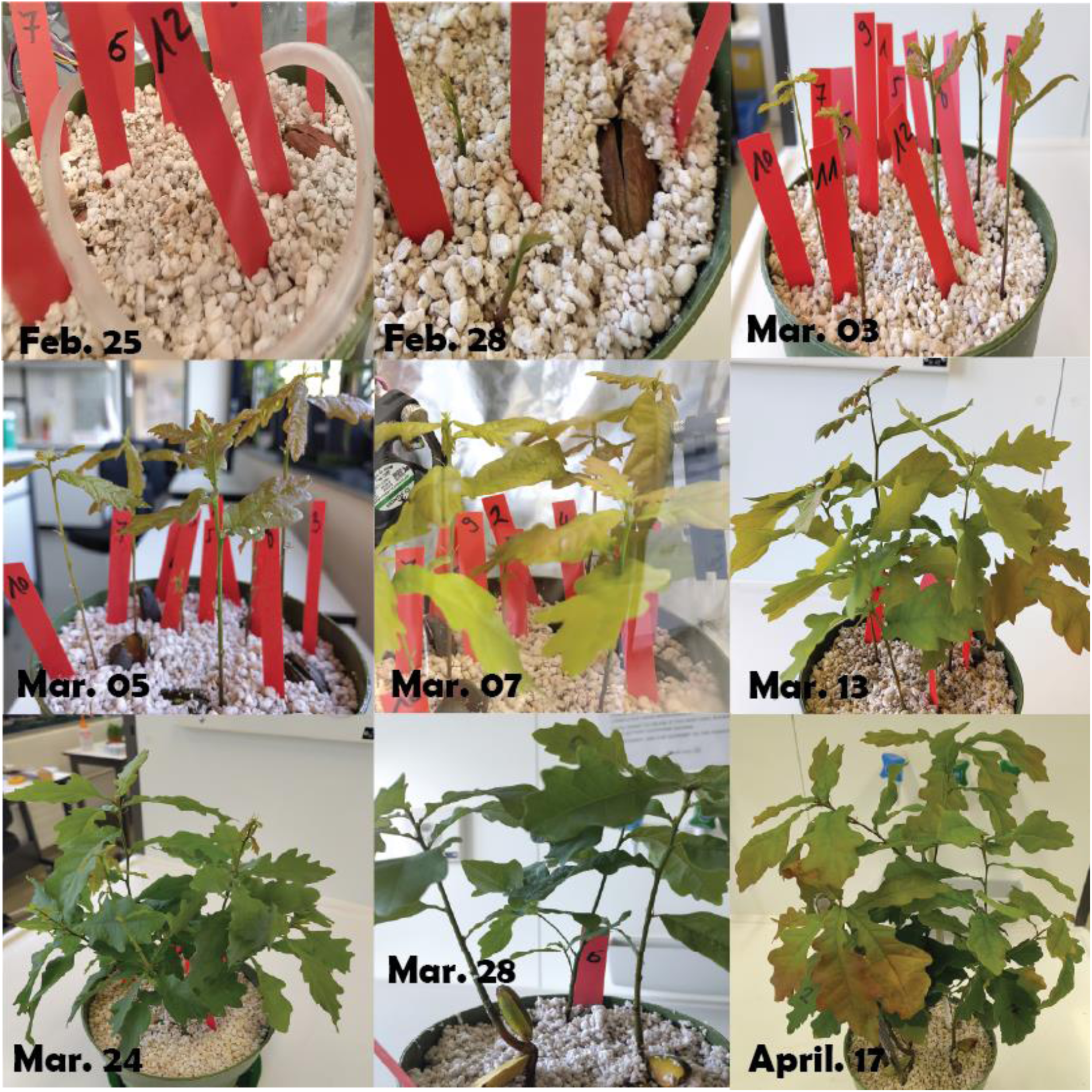
Time-lapse of the germinating *Quercus robur* acorns from the 25^th^ of February 2025 (3 days after sowing) until the 17th of April 2025, the day after the second round of temperature response curves.

Early in the germination process, net CO_2_ flux was consistently negative, indicating a fully heterotrophic metabolism (cf. Fig. 5, Feb. 25). This also highlights the system’s sensitivity to below-ground respiration. Daytime fluxes were more negative than nighttime, likely due to ∼12°C higher daytime temperature. As root and shoot growth progressed, CO_2_ flux remained negative – even after initial leaf emergence on March 3^rd^, 5^th^, and 7^th^ (Fig. 5), likely due to high growth respiration. A sharp nighttime peak in negative flux developed at the onset of darkness, likely reflecting high tissue respiration at still high temperature, while a small positive peak appeared at lights-on, possibly due to cooler morning temperatures. Increasing photosynthesis led daytime net CO_2_ flux turning positive on March 8^th^.

Cumulative CO_2_ (Fig. 4b) and C (Fig. 4c) balances reached a turning point between March 8^th^- and 10^th^, after which it turned from negative to positive. After this, the strongly increasing photosynthetic activity (Fig. 4a) led to a progressively increasing positive net CO_2_ flux during the daytime hours, while nighttime respiration remained did not increase at a similar rate. This led to a steep increase in cumulative CO_2_ (Fig. 4b) and C (Fig. 4c) balance. By March 20^th^, seedling growth finished and net CO_2_ flux plateaued, a mild drought was induced. This did not impact the early morning peak of net CO_2_ uptake (Fig. 4a) but decreased the net CO_2_ uptake during the later hours of the day. This resulting midday depression of photosynthesis resembles field observations (Muraoka et al., 2000; Koyama and Takemoto, 2014), though here it likely results from soil water limitation rather than photoinhibition.

Finally, a comparison of cumulative C balance with dry biomass (Fig. 4C) showed strong agreement: the calculated C gain per seedling was 609 mg glucose equivalent, closely matching the 544 mg average of seedlings harvested on March 24. The slight offset likely reflects developmental variability (Fig. 5) and sample size (i.e., only three per time point).

### Experiment 4: Shifts in whole-plant C balance with rising temperatures

The net CO_2_ flux responded strongly to increasing T_air_ (Figs. 6, S4), with notable differences between the two experimental rounds. Between the two rounds, total leaf area increased by 595 cm^2^ (from 873 cm^2^ to 1468 cm^2^), and leaf mass fraction (the percentage the leaf weight contributes to the total weight) rose from 15.7% to 34.4%. On the first day (average daytime T_air_ of 27.6°C for both rounds), net CO_2_ uptake was 4.4 times higher in the second round (1.15 µmol s^-1^) than in the first (0.26 µmol s^-1^). On the second day, daytime T_air_ averaged 33.6°C in the first round and 31.8°C in the second. Correspondingly, the net CO_2_ flux declined by 31% in the first round and 10% in the second (Fig. 6b). Despite this, flux on the second round (1.03 µmol s^-1^) remained 5.7 times higher than in the first (0.18 µmol s^-1^). On the third day, with average T_air_ reaching 42.9°C and 42.3°C respectively, the net CO_2_ flux dropped sharply in both rounds: to 14% of the original value in the first (0.04 µmol s^-1^) and to 30% in the second (0.35 µmol s^-1^), which was still 8.8 times higher than in the first.

**Figure 6:**
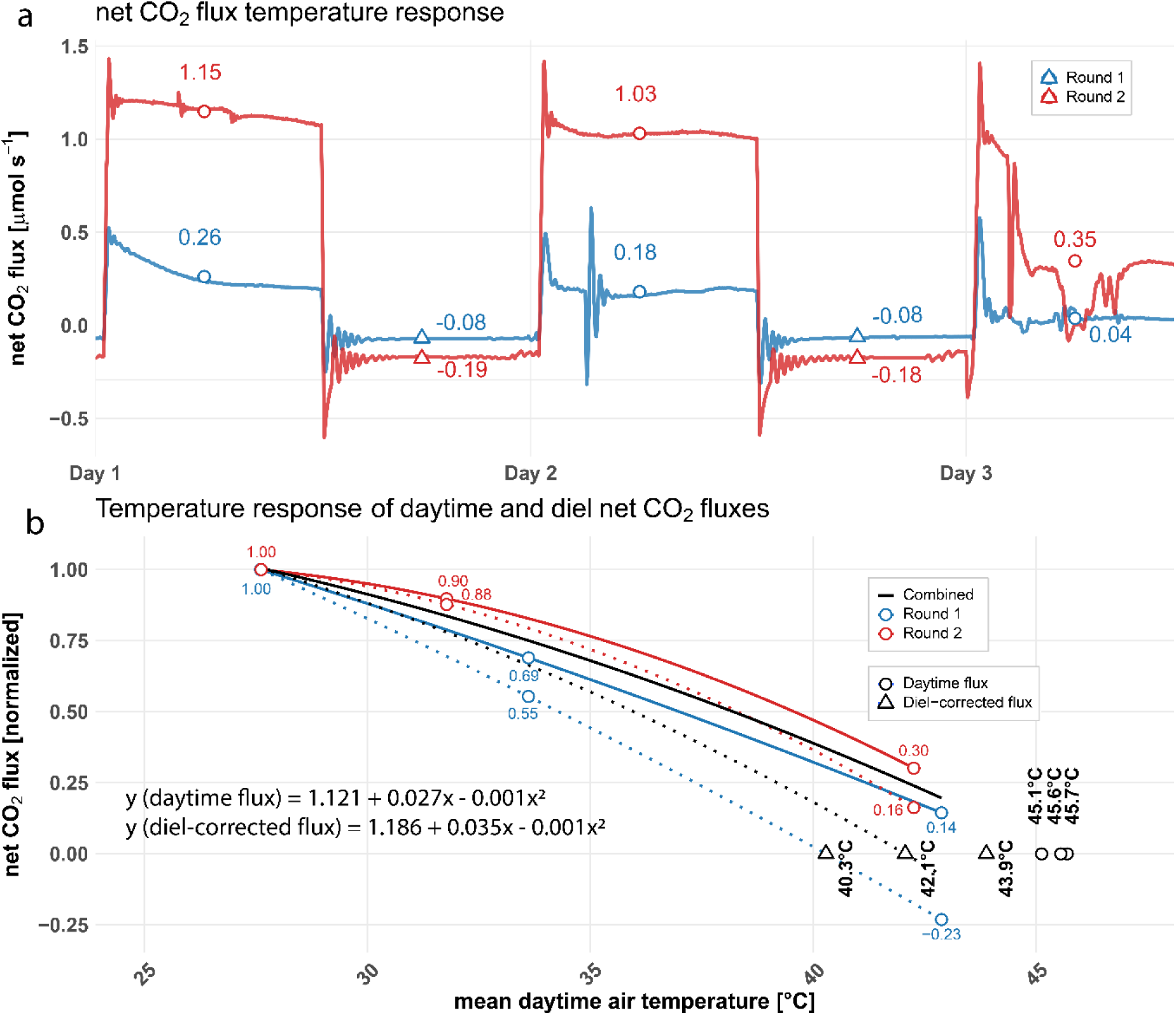
(a) The running means of the net CO_2_ flux in µmol s^-1^ during the three days with increasing air temperature (T_air_) of *Quercus robur* seedlings (all three seedlings combined) after their first (Round 1, blue, average daytime T_air_ 27.6, 33.6, and 42.9°C) and second (Round 2, red, average daytime T_air_ 27.6, 31.8, and 42.3°C) development stages finished, where negative values indicating a net CO_2_ release whereas positive values indicate a net CO_2_ uptake. Empty circles indicate the average net CO_2_ flux during daytime, while triangles indicate average nighttime flux at room temperature (∼22°C). (b) The corresponding temperature response (blue = round 1, red = round 2, black = combined) of the mean daytime net CO_2_ flux (straight line) and the diel-corrected net CO_2_ flux (i.e., average daytime flux – average nighttime flux; dashed line) normalized to the mean of the first day with a 27.6°C, the quadratic fit for the mean of both rounds, as well as the temperature thresholds at which the net CO_2_ flux turns negative (triangles for the diel-corrected, i.e. daytime – nighttime flux and circles for the daytime fluxes, number). The red and blue numbers indicate average values of the corresponding days and nights, respectively.

Quadratic regressions models fitted to the normalized mean daytime net CO_2_ flux (Fig. 6b) predicted the temperature at which net flux becomes zero to be 45.1°C (first round), 45.7°C (second round), and 45.6°C (combined data). When diel CO_2_ flux was calculated by subtracting nighttime from daytime values (Fig. 6b), the flux turned neutral at 40.3°C, 43.9°C, and 42.1°C for the first, second, and combined rounds, respectively.

However, these measurements were based on a single pseudo-replicate, and the temperature responses therefore overfitted. As such, these results should only be viewed as a proof of concept, illustrating the capabilities of this method with the PANDA system. Future experiments with appropriate replication are needed to robustly quantify how the net CO₂ flux responds to increasing temperature, and how this response is influenced by factors like plant size and leaf mass fraction.

## Conclusions

We presented a replicable, low-cost chamber system – PANDA - that enables accurate, continuous, and reliable monitoring of whole-plant net CO₂ flux. This study demonstrates that the PANDA system provides a practical and flexible platform for studying plant C balance at the whole-plant level in a laboratory setting. Its application includes (1) monitoring diel dynamics of net CO_2_ fluxes in both C_3_ and CAM plants, (2) investigating circadian rhythms under prolonged exposure to light or dark conditions, (3) tracking the C balance during germination and early growth, and (4) assessing responses to environmental stressors such as drought and high temperature. PANDA’s modular design allows for easy adaptation to diverse experimental needs, such as scaling chamber size, integrating active temperature control, or situating the entire setup within a climate-controlled environment.

We highlight the value of shifting from measurements isolated plant organs to whole-plant assessments, offering a more integrated perspective on plant C dynamics under changing environmental conditions. Given its simplicity, affordability, and reproducibility, the PANDA system holds strong potential to become a widely adopted tool for advancing our understanding of whole-plant responses to climate-related stresses, thereby contributing to improved predictions of plant behavior in future climates.

## Author contributions

PS conceptualized the chamber and planned the experiments. PF and PS built the chamber system. PS, PP and YL conducted the experimental work. PS led the writing of the manuscript. YL, PP, PF, MHL, YLZ, and CG critically revised the manuscript.

## Data availability

After publication, all raw data will be published on a publicly available respiratory (www.envidat.ch).

## References

Atkin, O.K., Edwards, E.J., Loveys, B.R., 2000. Response of root respiration to changes in temperature and its relevance to global warming. New Phytologist 147, 141–154. 10.1046/j.1469-8137.2000.00683.x

Ávila, E., Herrera, A., Tezara, W., 2014. Contribution of stem CO_2_ fixation to whole-plant carbon balance in nonsucculent species. Photosynt. 52, 3–15. 10.1007/s11099-014-0004-2

Ávila-Lovera, E., Haro, R., Choudhary, M., Acosta-Rangel, A., Pratt, R.B., Santiago, L.S., 2024. The benefits of woody plant stem photosynthesis extend to hydraulic function and drought survival in *Parkinsonia florida*. Tree Physiology 44, tpae013. 10.1093/treephys/tpae013

Ben-Haj-Salah, H., Tardieu, F., 1995. Temperature Affects Expansion Rate of Maize Leaves without Change in Spatial Distribution of Cell Length (Analysis of the Coordination between Cell Division and Cell Expansion). Plant Physiol. 109, 861–870. 10.1104/pp.109.3.861

Birami, B., Gattmann, M., Heyer, A.G., Grote, R., Arneth, A., Ruehr, N.K., 2018. Heat Waves Alter Carbon Allocation and Increase Mortality of Aleppo Pine Under Dry Conditions. Front. For. Glob. Change 1, 8. 10.3389/ffgc.2018.00008

Brauner, K., Birami, B., Brauner, H.A., Heyer, A.G., 2018. Diurnal periodicity of assimilate transport shapes resource allocation and whole-plant carbon balance. The Plant Journal 94, 776–789. 10.1111/tpj.13898

Brauner, K., Hörmiller, I., Nägele, T., Heyer, A.G., 2014. Exaggerated root respiration accounts for growth retardation in a starchless mutant of *A rabidopsis thaliana*. The Plant Journal 79, 82–91. 10.1111/tpj.12555

Campioli, M., Malhi, Y., Vicca, S., Luyssaert, S., Papale, D., Peñuelas, J., Reichstein, M., Migliavacca, M., Arain, M.A., Janssens, I.A., 2016. Evaluating the convergence between eddy-covariance and biometric methods for assessing carbon budgets of forests. Nat Commun 7, 13717. 10.1038/ncomms13717

Collins, A.D., Ryan, M.G., Adams, H.D., Dickman, L.T., Garcia-Forner, N., Grossiord, C., Powers, H.H., Sevanto, S., McDowell, N.G., 2021. Foliar respiration is related to photosynthetic, growth and carbohydrate response to experimental drought and elevated temperature. Plant Cell & Environment 44, 3853–3865. 10.1111/pce.14183

Colombi, T., Chakrawal, A., Herrmann, A.M., 2021. Carbon supply–consumption balance in plant roots: effects of carbon use efficiency and root anatomical plasticity. New Phytologist 233, 1542–1547. CSIRO - Australian Tropical Rainforest Plants, 2020. Online edition.

DaMatta, F.M., Avila, R.T., Cardoso, A.A., Martins, S.C.V., Ramalho, J.C., 2018. Physiological and Agronomic Performance of the Coffee Crop in the Context of Climate Change and Global Warming: A Review. J. Agric. Food Chem. 66, 5264–5274. 10.1021/acs.jafc.7b04537

Diao, H., Wang, A., Yuan, F., Guan, D., Dai, G., Wu, J., 2020. Environmental Effects on Carbon Isotope Discrimination from Assimilation to Respiration in a Coniferous and Broad-Leaved Mixed Forest of Northeast China. Forests 11, 1156. 10.3390/f11111156

Dutton, R.G., Jiao, J., Tsujita, M.J., Grodzinski, B., 1988. Whole Plant CO_2_ Exchange Measurements for Nondestructive Estimation of Growth. Plant Physiol. 86, 355–358. 10.1104/pp.86.2.355

Frantz, J.M., 2004. Night Temperature has a Minimal Effect on Respiration and Growth in Rapidly Growing Plants. Annals of Botany 94, 155–166. 10.1093/aob/mch122

Gansert, D., Burgdorf, M., 2005. Effects of xylem sap flow on carbon dioxide efflux from stems of birch (Betula pendula Roth). Flora - Morphology, Distribution, Functional Ecology of Plants 200, 444–455. 10.1016/j.flora.2004.12.005

Gessler, A., Roy, J., Kayler, Z., Ferrio, J.P., Alday, J.G., Bahn, M., Del Castillo, J., Devidal, S., García-Muñoz, S., Landais, D., Martín-Gomez, P., Milcu, A., Piel, C., Pirhofer-Walzl, K., Galiano, L., Schaub, M., Haeni, M., Ravel, O., Salekin, S., Tissue, D.T., Tjoelker, M.G., Voltas, J., Hoch, G., Resco De Dios, V., 2017. Night and day – Circadian regulation of night-time dark respiration and light-enhanced dark respiration in plant leaves and canopies. Environmental and Experimental Botany 137, 14–25. 10.1016/j.envexpbot.2017.01.014

Grossiord, C., Mareschal, L., Epron, D., 2012. Transpiration alters the contribution of autotrophic and heterotrophic components of soil CO_2_ efflux. New Phytologist 194, 647–653. 10.1111/j.1469-8137.2012.04102.x

Han, F., Wang, X., Zhou, H., Li, Y., Hu, D., 2017. Temporal dynamics and vertical variations in stem CO_2_ efflux of Styphnolobium japonicum. J Plant Res 130, 845–858. 10.1007/s10265-017-0951-3

Hand, D.W., 1973. A null balance method for measuring crop photo-synthesis in an airtight daylit controlled-environment cabinet. Agricultural Meteorology 12, 259–270. 10.1016/0002-1571(73)90024-1

Harris, R., 1992. Root-Shoot Ratios. AUF 18, 39–42. 10.48044/jauf.1992.009

Jackson, D.K., 1974. Some characteristics of perlite as an experimental growth medium. Plant Soil 40, 161–167. 10.1007/BF00011418

Kölling, K., George, G.M., Künzli, R., Flütsch, P., Zeeman, S.C., 2015. A whole-plant chamber system for parallel gas exchange measurements of Arabidopsis and other herbaceous species. Plant Methods 11, 48. 10.1186/s13007-015-0089-z

Koyama, K., Takemoto, S., 2014. Morning reduction of photosynthetic capacity before midday depression. Sci Rep 4, 4389. 10.1038/srep04389

Kufa, T., Burkhardt, M.J., 2011. Plant composition and growth of wild Coffea arabica: Implications for management and conservation of natural forest resources. International Journal of Biodiversity and Conservation 3, 131–141.

Kumar, D., Tieszen, L.L., 1980. Photosynthesis in *Coffea arabica*. I. Effects of Light and Temperature. Ex. Agric. 16, 13–19. 10.1017/S0014479700010656

Kuzyakov, Ya.V., Larionova, A.A., 2006. Contribution of rhizomicrobial and root respiration to the CO_2_ emission from soil (A review). Eurasian Soil Sc. 39, 753– 764. 10.1134/S106422930607009X

Lai, Z., Lu, S., Zhang, Y., Wu, B., Qin, S., Feng, W., Liu, J., Fa, K., 2016. Diel patterns of fine root respiration in a dryland shrub, measured in situ over different phenological stages. Journal of Forest Research 21, 31–42. 10.1007/s10310-015-0511-4

Lambers, H., Atkin, O.K., Millenaar, F.F., 2002. Respiratory Patterns in Roots in Relation to Their Functioning, in: Plant Roots. p. 57.

Lüttge, U., Hertel, B., 2009. Diurnal and annual rhythms in trees. Trees 23, 683–700. 10.1007/s00468-009-0324-1

Marler, T.E., Lindström, A.J., 2020. Diel patterns of stem CO_2_ efflux vary among cycads, arborescent monocots, and woody eudicots and gymnosperms. Plant Signaling & Behavior 15, 1732661. 10.1080/15592324.2020.1732661

Meehl, G.A., Tebaldi, C., 2004. More Intense, More Frequent, and Longer Lasting Heat Waves in the 21st Century. Science 305, 994–997. 10.1126/science.1098704

Muraoka, H., Tang, Y., Terashima, I., Koizumi, H., Washitani, I., 2000. Contributions of diffusional limitation, photoinhibition and photorespiration to midday depression of photosynthesis in *Arisaema heterophyllum* in natural high light. Plant Cell & Environment 23, 235–250. 10.1046/j.1365-3040.2000.00547.x

Nunes, M.A., Bierhuizen, J.F., Ploegman, C., 1968. STUDIES ON PRODUCTIVITY OF COFFEE: I. EFFECT OF LIGHT, TEMPERATURE AND CO_2_ CONCENTRATION ON PHOTOSYNTHESIS OF COFFEA ARABICA. Acta Botanica Neerlandica 17, 93–102. 10.1111/j.1438-8677.1968.tb00109.x

Osmond, C.B., 1978. Crassulacean Acid Metabolism: A Curiosity in Context. Annu. Rev. Plant. Physiol. 29, 379–414. 10.1146/annurev.pp.29.060178.002115

Osmond, C.B., Nott, D.L., Firth, P.M., 1979. Carbon assimilation patterns and growth of the introduced CAM plant Opuntia inermis in Eastern Australia. Oecologia 40, 331–350. 10.1007/BF00345329

Patono, D.L., Eloi Alcatrāo, L., Dicembrini, E., Ivaldi, G., Ricauda Aimonino, D., Lovisolo, C., 2023. Technical advances for measurement of gas exchange at the whole plant level: Design solutions and prototype tests to carry out shoot and rootzone analyses in plants of different sizes. Plant Science 326, 111505. 10.1016/j.plantsci.2022.111505

Perkins-Kirkpatrick, S.E., Lewis, S.C., 2020. Increasing trends in regional heatwaves. Nat Commun 11, 3357. 10.1038/s41467-020-16970-7

Perry, T.O., 1982. The Ecology of Tree Roots and the Practical Significance Thereof. isa 8, 197–211. 10.48044/jauf.1982.047

Prinn, R.G., 2013. Development and application of earth system models. Proc. Natl. Acad. Sci. U.S.A. 110, 3673–3680. 10.1073/pnas.1107470109

Resco De Dios, V., Gessler, A., Ferrio, J.P., Alday, J.G., Bahn, M., Del Castillo, J., Devidal, S., García-Muñoz, S., Kayler, Z., Landais, D., Martín-Gómez, P., Milcu, A., Piel, C., Pirhofer-Walzl, K., Ravel, O., Salekin, S., Tissue, D.T., Tjoelker, M.G., Voltas, J., Roy, J., 2016. Circadian rhythms have significant effects on leaf-to-canopy scale gas exchange under field conditions. Gigascience 5, s13742–016-0149-y. 10.1186/s13742-016-0149-y

Salvatori, N., Giorgio, A., Muller, O., Rascher, U., Peressotti, A., 2021. A low-cost automated growth chamber system for continuous measurements of gas exchange at canopy scale in dynamic conditions. Plant Methods 17, 69. 10.1186/s13007-021-00772-z

Schneider, C.A., Rasband, W.S., Eliceiri, K.W., 2012. NIH Image to ImageJ: 25 years of image analysis. Nat Methods 9, 671–675. 10.1038/nmeth.2089

Sevanto, S., Mcdowell, N.G., Dickman, L.T., Pangle, R., Pockman, W.T., 2014. How do trees die? A test of the hydraulic failure and carbon starvation hypotheses. Plant Cell & Environment 37, 153–161. 10.1111/pce.12141

Surridge, C., 2019. Uncovering cryptic CAM. Nature Plants 5, 3–3. 10.1038/s41477-018-0351-2

Teskey, R.O., Saveyn, A., Steppe, K., McGuire, M.A., 2008. Origin, fate and significance of CO_2_ in tree stems. New Phytologist 177, 17–32. 10.1111/j.1469-8137.2007.02286.x

Tomlinson, P.B., 1990. Palm stem—mechanics, age determination, and hydraulics, in: Tomlinson, P.B. (Ed.), The Structural Biology of Palms. Oxford University Press, p. 0. 10.1093/oso/9780198545729.003.0007

Trumbore, S.E., Angert, A., Kunert, N., Muhr, J., Chambers, J.Q., 2013. What’s the flux? Unraveling how CO_2_ fluxes from trees reflect underlying physiological processes. New Phytologist 197, 353–355. 10.1111/nph.12065

Wang, W., Hoch, G., 2022. Negative effects of low root temperatures on water and carbon relations in temperate tree seedlings assessed by dual isotopic labelling. Tree Physiology 42, 1311–1324. 10.1093/treephys/tpac005

Wang, W., Wang, H., Zu, Y., Li, X., Koike, T., 2006. Characteristics of the temperature coefficient, Q 10, for the respiration of non-photosynthetic organs and soils of forest ecosystems. Front. Forest. China 1, 125–135. 10.1007/s11461-006-0018-4

Weber, R., Schwendener, A., Schmid, S., Lambert, S., Wiley, E., Landhäusser, S.M., Hartmann, H., Hoch, G., 2018. Living on next to nothing: tree seedlings can survive weeks with very low carbohydrate concentrations. New Phytologist 218, 107–118. 10.1111/nph.14987

Werner, C., Fasbender, L., Romek, K.M., Yáñez-Serrano, A.M., Kreuzwieser, J., 2020. Heat Waves Change Plant Carbon Allocation Among Primary and Secondary Metabolism Altering CO_2_ Assimilation, Respiration, and VOC Emissions. Front. Plant Sci. 11, 1242. 10.3389/fpls.2020.01242

Winter, K., Holtum, J.A.M., 2015. Cryptic crassulacean acid metabolism (CAM) in Jatropha curcas. Functional Plant Biology 42, 711–717.

Winter, K., Sage, R.F., Edwards, E.J., Virgo, A., Holtum, J.A.M., 2019. Facultative crassulacean acid metabolism in a C_3_–C4 intermediate. Journal of Experimental Botany 70, 6571–6579. 10.1093/jxb/erz085

Zhang, H., Wang, H., Wright, I.J., Prentice, I.C., Harrison, S.P., Smith, N.G., Westerband, A.C., Rowland, L., Plavcová, L., Morris, H., Reich, P.B., Jansen, S., Keenan, T., Nguyen, N., 2025. Thermal acclimation of stem respiration implies a weaker carbon-climate feedback.

Zhao, J., Hartmann, H., Trumbore, S., Ziegler, W., 2013. High temperature causes negative whole-plant carbon balance under mild drought. New Phytologist 200, 330–339. 10.1111/nph.12400

